# MK-801-induced behavioral and dopaminergic responses in the shell part of the nucleus accumbens in adult rats are disrupted after neonatal blockade of the ventral subiculum

**DOI:** 10.1101/2021.02.16.431488

**Authors:** Hana Saoud, Elora Kereselidze, Séverine Eybrard, Alain Louilot

**Affiliations:** University of Strasbourg, INSERM U 1114, Faculty of Medicine, FMTS, Strasbourg, France

**Keywords:** MK-801 administration, animal modeling, schizophrenia, TTX postnatal inactivation, dopamine, *in vivo* voltammetry

## Abstract

The present study was conducted in the context of animal modeling of schizophrenia. It investigated in adult rats, after transient neonatal blockade of the ventral subiculum (VSub), the impact of a very specific non-competitive antagonist of NMDA receptors (MK-801) on locomotor activity and dopaminergic (DAergic) responses in the dorsomedial *shell* part of the nucleus accumbens (Nacc), a striatal subregion described as the common target region for antipsychotics.

The functional neonatal inactivation of the VSub was achieved by local microinjection of tetrodotoxin (TTX) at postnatal day 8 (PND8). Control pups were microinjected with the solvent phosphate buffered saline (PBS). Locomotor responses and DAergic variations in the dorsomedial *shell* part of the Nacc were measured simultaneously using *in vivo* voltammetry in awake, freely moving animals after sc administration of MK-801. The following results were obtained: 1) a dose-dependent increase in locomotor activity in PBS and TTX animals, greater in TTX rats/PBS rats; and 2) divergent DAergic responses for PBS and TTX animals. A decrease in DA levels with a return to around basal values was observed in PBS animals. An increase in DA levels was obtained in TTX animals. The present data suggest that neonatal blockade of the VSub results in disruption in NMDA glutamatergic transmission, causing a disturbance in DA release in the dorsomedial *shell* in adults rats. In the context of animal modeling of schizophrenia using the same approach it would be interesting to investigate possible changes in postsynaptic NMDA receptors-related proteins in the dorsomedial *shell* region in the Nacc.

## 1. Introduction

The present study concerns animal modeling of the pathophysiology of schizophrenia. Such modeling appears crucial for gaining a better understanding of disease and ultimately for developing new treatments, given that existing treatments are not completely satisfactory (Leucht et al., 2017).

Schizophrenia is a complex heterogeneous mental disorder, beginning in adolescents and young people, and whose etiology and pathophysiology are not fully understood (Tandon et al., 2013). Different hypotheses have been proposed to explain this particular pathophysiology, the oldest being the dopaminergic (DAergic) hypothesis first put forward in 1966 (Van Rossum, 1966; see also Seeman, 1987). According to this hypothesis, which is still one of the best established and best characterized, there exists in schizophrenia a striatal hyperdopaminergy and a prefrontal cortical hypodopaminergy. The hypothesis of a striatal hyperdopaminergy is reinforced by data that show that antipsychotics all exhibit the property to block DAergic D2 receptors to some degree (Kaar et al., 2020). As regards striatal hyperdopaminergic activity, meta-analyses of brain imaging studies suggest an increase in DA release in either both the dorsal and ventral parts of the striatum (Nikolaus et al., 2019) or only in the dorsal striatum (mainly in the associative striatum)(McCutcheon et al., 2018). However, subdivisions of the ventral striatum and differential changes in these sub-territories have never be reported in such meta-analyses, whereas preclinical studies support the view that the *shell* subdivision of the nucleus accumbens (Nacc), part of the ventral striatum, is the shared anatomical target for all antipsychotics (typical and atypical antipsychotics), with the dorsolateral striatum being the preferential target for typical antipsychotics (Deutch et al., 1992, Merchant and Dorsa, 1993, Mo et al., 2005; Jennings et al., 2006; Natesan et al., 2006; Collins et al., 2014).

A second hypothesis, the glutamatergic hypothesis of schizophrenia, was proposed in parallel following observations showing the emergence of psychotic symptoms in healthy subjects and the worsening of such symptoms in patients with schizophrenia following administration of N-methyl-D-aspartate receptor antagonists (NMDA), with glutamate being the main agonist of these receptors. In other words, there are extensive data to suggest that in patients with schizophrenia there is a dysfunction in glutamatergic transmission involving NMDA receptors (Itil et al., 1967; Coyle, 2012). Interestingly, postmortem studies have recently reported reduced postsynaptic density (PSD)size at excitatory synapses in the Nacc (*core* and *shell* subregions) of patients with schizophrenia compared to normal controls. This reduction could be due to a disruption of the glutamatergic transmission in the Nacc in patients (McCollum et al., 2015).

More recently, some 30 years ago, a third hypothesis was put forward according to which schizophrenia is the result of disconnections in the brain between different integrative regions and, at least in certain cases, of neurodevelopmental origin. Anomalies during early neurodevelopment, might give rise to connection disturbances involving abnormal myelination, and/or aberrant synaptic connectivity. According to this hypothesis, some brain regions, such as the parahippocampal/hippocampal formation, prefrontal cortex, and striatal regions, would be more specifically affected by these brain disconnections (Friston, 1996; McGlashan and Hoffman, 2000; Talamini et al., 2005; Karlsgodt et al., 2008; Fornito et al., 2012; Giezendanner et al., 2013; Weinberger, 2017; see also Meyer and Louilot, 2014).

The dysfunctioning of DAergic and glutamatergic neurotransmissions in schizophrenia could be linked, with the hypothesis of cerebral disconnection between the different integrative regions or the brain (Meyer and Louilot, 2014) providing the explanation. In this context, it is important to note that the ventral subiculum (VSub), which is part of the hippocampal formation allowing the output of the signals from the CA1 region of the hippocampus, seems to be particularly sensitive to neurodevelopmental alterations in schizophrenia (Rioux et al., 2003; Moyer et al., 2015). Moreover, a reduction in the volume of the *left* VSub volume was observed in patients with first episode psychosis (Briend et al., 2020). In other respects regarding this conceptual framework, it is worth mentioning that anatomical projections from the VSub towards the *shell* subregion have been described, as well as NMDA-type glutamatergic receptors present at the presynaptic level on the DAergic terminals in this part of the Nacc (Meredith and Totterdell, 1999). It is thus possible to consider that disconnection of the VSub might have consequences for DA release in the *shell* of the Nacc through NMDA receptors.

In light of all the above-mentioned data, the aim of the present study conducted on animals (rats) was to determine the impact at adulthood of neonatal functional blockade of the left VSub on locomotor activity and DAergic responses in the left dorsomedial *shell* part of the Nacc after the adults had been administered the very specific non-competitive NMDA receptor antagonist MK-801 (or dizocilpine) (Reynolds et al., 1987; Ogden and Traynelis, 2011). Locomotor hyperactivity is widely considered to be a valid behavioral marker in the context of animal modeling of the pathophysiology of schizophrenia, due to parallels that can be drawn between the locomotor hyperactivity observed in rodents and the emergence of, or increase in the positive symptoms described in humans following the administration of psychoactive drugs (Van den Buuse, 2010).

DA levels in the dorsomedial *shell* are measured using *in vivo* voltammetry in adult animals (11 weeks old) in chronic preparation and free to move. Neonatal functional blocking of the VSub is obtained by local microinjection of TTX on postnatal day 8 (PND8). TTX injected at a critical point in the neurodevelopmental period in rats (equivalent to the end of the second trimester of pregnancy in humans) gives rise to a decrease in myelination, a disruption of synaptic refinement and dendritic maturation, as well an increase in NMDA receptor traffic, all of which are described in patients with schizophrenia (see Tagliabue et al., 2017 for discussion).

## 2. MATERIAL AND METHODS

### 2.1. Animals and experimental design

Experiments were performed on Sprague Dawley rats issued from gestant mothers from the breeding center Janvier-Labs (Le Genest St-Isle, France). All the experiments comply with the ARRIVE guidelines and were carried out in accordance with Directive 2010/63/EU on the protection of animals used for scientific purposes. The experimental procedures were all approved by the Strasbourg Regional Ethics Committee on Animal Experimentation (CREMEAS-CEEA35) and the French Ministry of Higher Education, Research and Innovation (authorization APAFIS# 1553-2015082613521475 v3). All the animals used for the experiment were kept throughout under standard controlled temperature conditions (22±2° C) and had *ad libitum* access to food and water. Every effort was made to minimize the number of animals used and their suffering. During gestation, mothers were kept in individual Plexiglas cages on a 12 hr light/dark cycle (lights on at 7 am). The day pups were born was defined as postnatal day 0 (PND0). On PND8, the neonates were randomly given a microinjection of either PBS (control group) or TTX (experimental group) in the left ventral subiculum (VSub). After weaning at postnatal day PND21, only male rats were used. On PND56, male rats were acclimatized to a reversed light/dark cycle (lights off from 11 am to 11 pm) before being fitted on PND70 with a specially designed microsystem that allowed parallel measurement of behavioral responses and dopamine changes (as measured with *in vivo* voltammetry) in freely moving adult rats. About one week after surgery (on PND77), behavioral and voltammetric recordings were made during the dark period of the reversed light/dark cycle.

### 2.2. Neonate surgery

On PND8, functional inactivation of the left VSub was carried out under general inhalation anaesthesia using isoflurane (IsoFlo®, Zoetis France, Malakoff, France) and stereotaxic surgery. The experimental procedure was identical to that previously reported in detail (Meyer et al., 2009; Meyer and Louilot, 2011, 2012; Usun et al., 2013; Saoud et al., 2020). According to pilot studies, naive unoperated animals did not differ from control rats microinfused with PBS. All the pups from each litter, both males and females, received microinjection of PBS and TTX (100 μM) on a random basis to avoid changes in the mother’s behavior towards one sex as opposed to the other (neglect or exacerbated attention regarding operated, or, non-operated rats) during the period prior to weaning on PND21. A volume of 0.3μl was infused over 2min 15s at coordinates 0.6 mm anterior to bregma (AP), 4.15 mm lateral to the midline (L), and 4.85 mm below the cortical surface (H). As presented and discussed in prior studies (Meyer et al., 2009; Meyer and Louilot, 2011, 2012; Usun et al., 2013) the microinfused TTX amount in the VSub corresponds to about 10 ng (100 μM × 0.3 μL), which appears to be sufficient to allow transient inactivation of the VSub for between 4 and 48hrs, within a functional inactivation radius of no more than 0.67 mm. PBS and TTX solutions were tinted with Evans Blue (Sigma, France), a dye which remains visible in the cerebral tissue and means the microinjection sites can be localized several weeks later. It was possible to identify neonates by means of their three-digit metal ear tags (ref. 52-4716, Harvard Apparatus, Les Ulis, France). After surgery, they were placed under a heating lamp until they woke up and then returned to their mothers.

### 2.3. Adult surgery

On PND70 a specially designed microsystem used to monitor the behavior and DAergic changes in freely moving animals simultaneously (Louilot et al., 1987) was stereotaxically implanted in the male rats having previously undergone neonatal microinjection in the VSub. The adult animals had the microsystem implanted in the left dorsomedial *shell* part of the Nacc with the following coordinates: +10.6 mm from the interaural line defined by the ear bars (AP coordinate); 1.6 mm from the midline defined by the sagittal suture (L coordinate); −7.1 mm from the surface of the cortex (H coordinate) (Paxinos and Watson, 2014). Adult rats, weighing 400 ± 25 g at the time of surgery, were anaesthetized by intraperitoneal injection of chloral hydrate (400 mg/kg; 8 ml/kg i.p.) (Field et al., 1993), then placed in a stereotaxic frame (M2E-Unimécanique, Eaubonne, France) with the incisor bar set at −3.3mm relative to the interaural line. To ensure as full analgesia as possible, an additional subcutaneous (s.c.) injection of a local anaesthetic (Lurocaïne®, Vetoquinol S.A, France) was carried out at the level of the animal’s head before surgery. After surgical procedure there was a period of at least 7 days so that the operated rats could recuperate before pharmacological studies.

### 2.4. Pharmacological studies

#### 2.4.1. Behavioral analysis

The behavioral index used in this study is locomotor activity. Changes in locomotor activity after MK-801 administration were investigated in a total of 54 implanted rats as described above. The animals were given about 1 hr to get use to the experimental cage (24 cm wide x 27 cm long x 44 cm high) before being injected on a random basis with either a s.c. injection of NaCl (0.9%) (control groups) or one of the two doses (0.1 mg/kg or 0.2 mg/kg) of MK-801 (experimental groups). Following the injection, locomotor activity was recorded for two further hours. During the pre-injection and post-injection periods, locomotor activity was measured simultaneously with the extracellular levels of DA in the dorsomedial *shell* part of the Nacc. Behavior was analyzed by direct observation after video recording. The floor of the experimental cage was split virtually into four identical zones. Each time the rat crossed over a virtual line separating two zones it counted as one crossing. Locomotor activity is expressed as mean ± SEM of the number of crossings per 10 min periods.

#### 2.4.2 DAergic variations: Voltammetric analysis

The extracellular levels of DA in the dorsomedial *shell* part of the Nacc were measured simultaneously with the locomotor activity using electrochemical procedures already described in detail (Meyer et al., 2009; Meyer and Louilot, 2011; 2012; Gonzalez-Mora et al., 2017). In short, selective detection of extracellular DA *in vivo* was provided by differential normal pulse voltammetry (DNPV) coupled to electrochemically pre-treated carbon fiber microelectrodes and computerized mathematical analysis of the DNPV signal (Gonzalez-Mora et al., 2017). For each animal, the average amplitude of the last 10 DA peaks recorded during the control period preceding the s.c. injection was calculated and constituted the 100 % value. The variations in the DA signal recorded per min in the *shell* part of the Nacc were then calculated as percentages (average ± SEM) of the average pre-injection value calculated on the basis of the last 10 DA peaks.

### 2.5. Statistics

The statistical significance of behavioral and DAergic changes was determined using multifactorial analysis of variance (ANOVA) with repeated measures for the time factor. Only between-subject ANOVAs are set out here unless stated otherwise. The independent variables were, on one hand, the postnatal microinjection, a two-level factor (PBS, TTX), and, on the other hand, the dose administered in adulthood, a three-level factor (NaCl 0.9%, MK 801 0.1 mg/kg or MK 801 0.2 mg/kg). Moreover, the dependent variables were the crossings (number) in the behavioral study as well as the changes in extracellular DA levels (%) in the voltammetric study. For the behavioral study, *post hoc* contrast analyses of the ANOVA supplemented the statistical analysis so as to test specific hypotheses (Rosenthal et al., 2000). Statistical results where p< 0.05 were considered significant.

### 2.6. Histology

In order to locate the voltammetric recording site, at the end of the experiment each animal underwent an electrocoagulation procedure already described in detail (Meyer et al., 2009). As previously mentioned (see above 2.2), the vital dye Evans Blue is added to PBS or TTX solutions as a way of identifying the postnatal microinjection site. Between 24 and 48 hrs after the experiments, the rats received a lethal injection of 5 ml/kg of the anaesthetic sodium pentobarbital (Doléthal®) before being rapidly intracardially perfused with 60 ml NaCl (0.9%) and 100 ml formalin (4%). After decapitation, animals brains were quickly removed and held at 4-8 °C in a solution comprising paraformaldehyde (PFA) (8%) and sucrose (30%) for at least 24 hours. After brain-sectioning, the histological sections were stained with Thionin (Nissl stain) for the electrocagulation site at the *shell* level in the Nacc or with Neutral Red to reveal the Evans Blue mark in the VSub region. The different structures were identified with precision by means of a dissecting microscope according to the Paxinos and Watson atlas (2014).

## 3. RESULTS

### 1. Histological results (Figure 1)

Typical sites for voltammetric recording in the *shell* of the left Nacc and for neonatal microinjection in the left VSub are identified *post mortem* on frontal sections of adult rats stained with thionine and neutral red respectively (Figure.1). The macroscopic qualitative analysis of the histological sections displays an absence of gliosis, and anatomical differences between the adult rats microinjected with the PBS solution and those microinjected with the TTX solution at PND8, at the level of both the shell part of the Nacc and the VSub.

**Figure 1:**
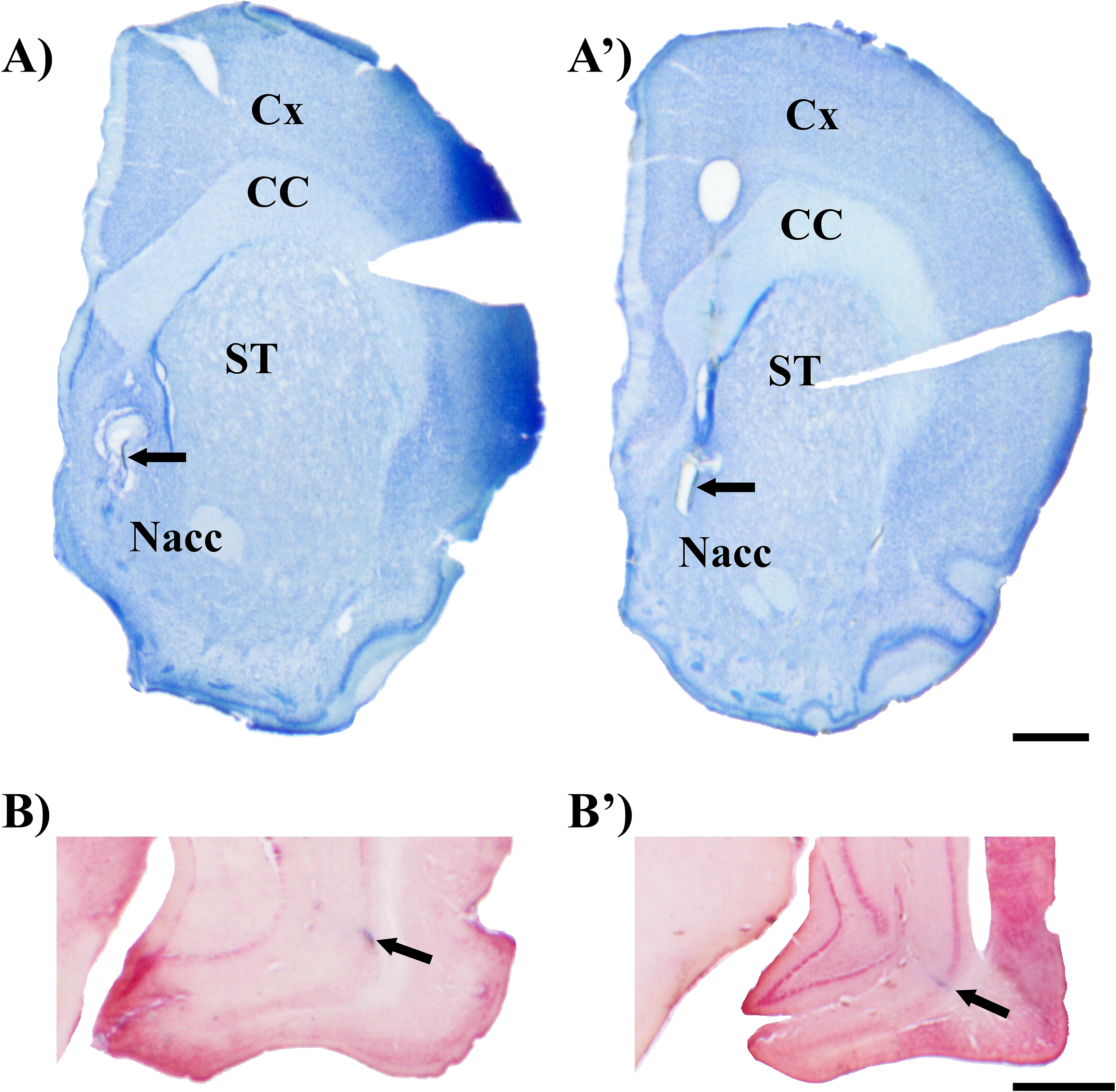
Microphotographs corresponding to typical recording sites in the left dorsomedial *shell* part of the accumbens nucleus (A and A’) and typical sites of neonatal microinjection in the left ventral subiculum (B and B’). The implantation site for the carbon fiber microelectrode in adult rats microinjected with the PBS (A) or TTX (A’) solution at PND8 is visualized after electrocoagulation carried out at the end of the recording and staining of the sections with Thionine; the fiber remained *in situ* at least in part (arrow). The site of neonatal microinjection of PBS (B) or TTX (B’) in the ventral subiculum (VSub) is identified by means of the trace left by Evans Blue (arrows) present in the PBS and TTX solutions microinjected at PND8 and revealed after coloring the brain sections with Neutral Red. Scale bar: 1 mm. Cx: cortex. CC: corpus callosum. ST: striatum. Nacc: nucleus accumbens.

After histological control, only animals in respect of which the neonatal microinjection site corresponded to the left VSub were considered for the whole study. Furthermore, rats presenting an implantation site of the carbon fiber microelectrode not clearly located in the dorsomedial part of the *shell* region of the left Nacc were excluded from the voltammetric analysis but included for the behavioral analysis. A total of 54 animals were used for behavioral analysis and 37 for voltammetric analysis

### 2. Behavioral results (Figure 2)

Before subcutaneous injection of NaCl 0.9%, MK-801 0.1 mg/kg or MK-801 0.2 mg/kg, the locomotor activity of the 54 animals split between the different groups was not significantly different (Figure 2). The general ANOVA performed on the 54 animals for the number of crossings during the 10 min preceding the sc injection shows no significant effect of the dose of MK-801 administered (F[2,48] = 2.94; ns), neither of the neonatal microinjection (F[1,48] = 0.28; ns), nor the dose x microinjection interaction (F[2,48] = 2.2; ns).

**Figure 2:**
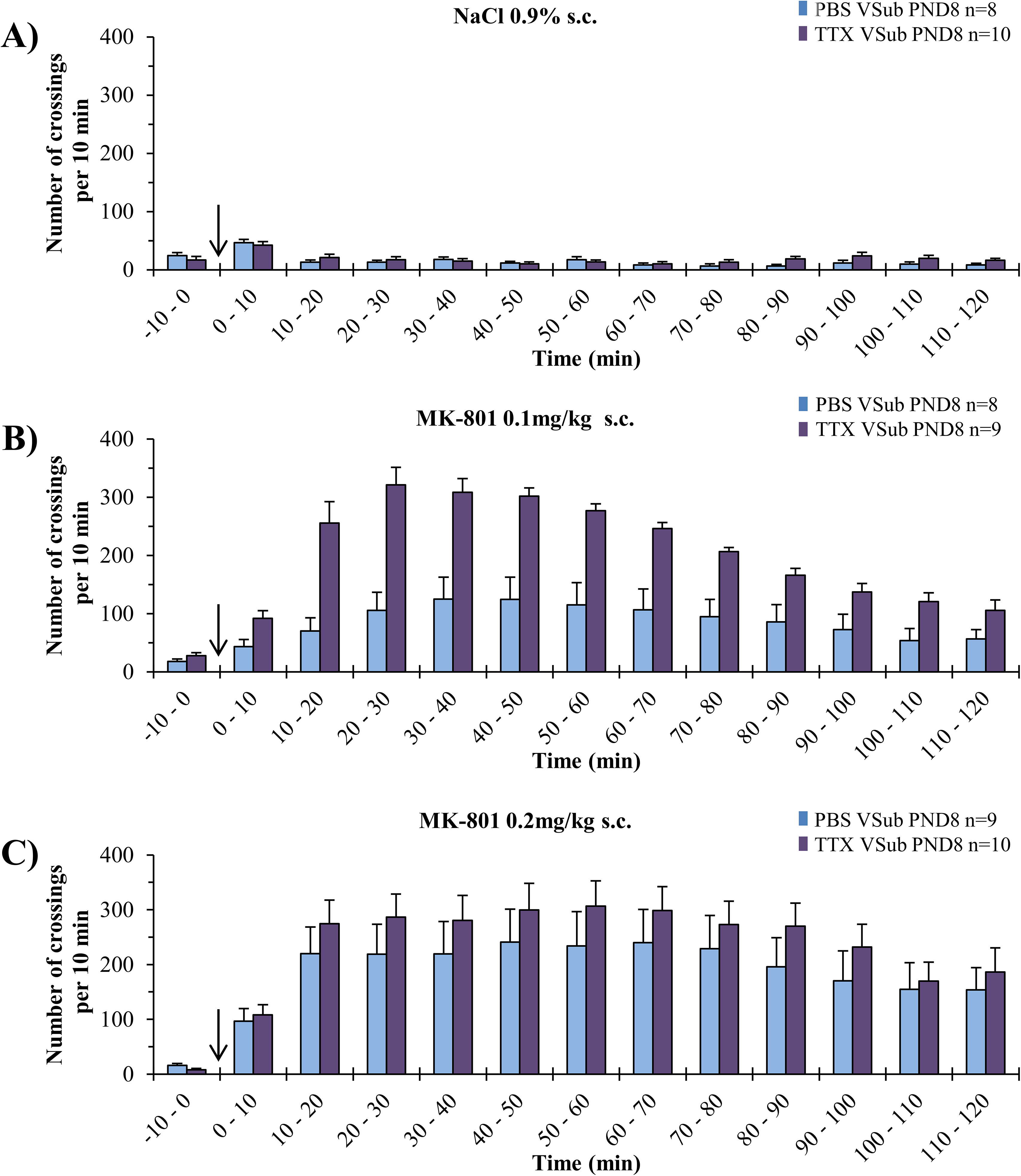
Locomotor activity after administration of MK-801 in adult rats having undergone neonatal inactivation of the left ventral subiculum (VSub) by microinjection of TTX at postnatal day 8 (PND8). Each adult animal micro-injected in the left VSub with the PBS (blue) or the TTX (purple) solution at PND8 receives, at random, a sc injection (arrow) of NaCl 0.9% (A), MK-801 0.1 mg/kg (B), or MK-801 0.2 mg/kg (C). Locomotor activity is measured as the number of crossings per 10 min period. The results are expressed as mean + SEM of the number of crossings per 10 min period. n denotes the number of rats per group. A factorial ANOVA was used for statistical analysis.

The general ANOVA performed on the 54 animals for the number of crossings during the 60 min post-injection period shows a significant effect for the administered dose (NaCl 0.9%; MK-801 0.1 mg/kg; MK-801 0.2 mg/kg) (F[2,48] = 24.72; p <0.00001), for neonatal microinjection (PBS/TTX) (F[1,48] = 7.83; p < 0.01), and for the dose x injection interaction (F [2,48] = 3.83; p <0.05). The general ANOVA performed on all 54 animals for the number of crossings during the 120 min post-injection period reveals a significant effect for the dose administered (NaCl 0.9%; MK-801 0.1 mg/kg; MK-801 0.2 mg/kg) (F [2,48] = 24.32; p <0.00001), and for neonatal microinjection (PBS/TTX) (F [1,48] = 5.7; p <0.05). On the other hand, there is no significant effect for the dose x injection interaction (F[2,48] = 1.9; n.s).

Contrast analysis of ANOVA was performed for the first 60 min post-injection to test the hypothesis that locomotor activity is significantly different depending on the neonatal microinjection (PBS or TTX) for the doses of MK-801 0.1 mg/kg and MK-801 0.2 mg/kg. Contrast analysis for the first 60 min post-injection reveals a significant effect for the neonatal microinjection factor for the dose 0.1 mg/kg sc (F[1,48] = 12.47; p <0.001) but not for the dose 0.2 mg/kg sc (F[1,48] = 1.58; ns).

As regards the time course of locomotor responses, the general ANOVA performed for the 54 animals during the 60 min post-injection period reveals a highly significant effect of the time factor (F[5,240] = 31.9; p <0.00001), a highly significant effect of the time x dose interaction (F[10,240] = 15.21; p <0.00001), a significant effect of the time x microinjection interaction (F[5,240] = 4.48; p <0.001), but no significant time x dose x microinjection interaction (F[10,240] = 2.31; ns). Concerning the 120 min post-injection, the ANOVA highlights significant effects for the time factor(F[11,528] = 27.1; p <0.00001), the time x dose interaction (F[22,528] = 10.74; p <0.00001), the time x microinjection interaction (F[11,528] = 3.68; p <0.0001), and the time x dose x interaction microinjection (F[22,528] = 2.31; p <0.001).

In summary, locomotor activity following administration of MK-801 is dependent on the dose (NaCl 0.9%; MK-801 0.1 mg/kg; MK-801 0.2 mg/kg) and neonatal microinjection in the left VSub(PBS or TTX). More specifically, the nature of the neonatal microinjection has a significantly different effect on the locomotor responses induced by MK-801 for the 0.1 mg/kg sc dose, but not the higher 0.2 mg/kg sc dose, during the 60 min post-injection period. In addition, the temporal evolution of the locomotor responses appears to depend on the dose of MK-801 administered as well as on neonatal microinjection for the 60 min as well as the 120 min post-injection.

### 3. DAergic Variations in the Nacc Shell (Figure 3)

The general ANOVA performed on the 37 animals for variations in the DAergic signal during the first 60 min post-injection shows a significant effect of neonatal microinjection (PBS or TTX) (F[1,31] = 7.04; p <0.05) but no significant dose effect (NaCl 0.9%; MK-801 0.1 mg/kg; MK-801 0.2 mg/kg) (F[2,31] = 0.34; ns), and no significant dose x microinjection interaction (F[2,31] = 1.37; ns).

**Figure 3:**
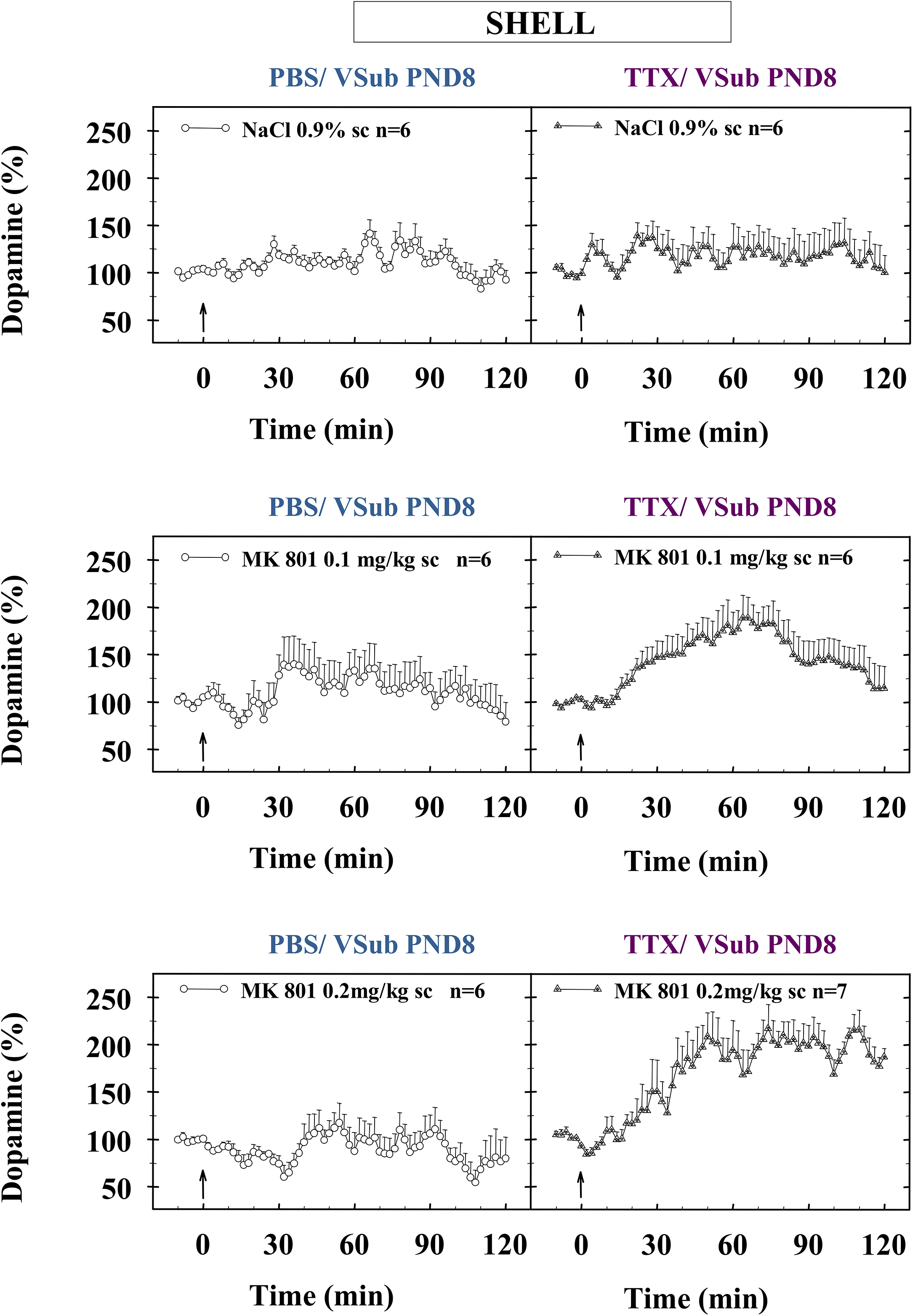
Dopaminergic variations within the dorsomedial *shell* part of the left nucleus accumbens after administration of MK-801 in adult rats having undergone neonatal functional inactivation of the VSub with TTX at PND8. Each adult animal microinjected in the left VSub with the PBS (ο) or TTX (∆) solution at PND8 received, at random, a sc injection (arrow) of NaCl 0.9% (A), MK-801 0.1 mg/kg (B), or MK-801 0.2 mg/kg (C). The extracellular levels of DA in the *shell* of the left Nacc were measured using Differential Normal Pulse Voltammetry (DNPV) with computer-assisted numerical analysis of the DNPV signal. The height of each peak is expressed as a percentage of the basal level (100%) corresponding to the mean amplitude of the last 10 peaks of DA recorded during the control period preceding the sc injection (arrow). Each point represents the mean + SEM of the percentage values obtained every 2 minutes. n corresponds to the number of rats per group. A factorial ANOVA was used for statistical analysis.

Regarding the 120 min post-injection, the performed ANOVA reveals a significant effect of neonatal microinjection (F[1,31] = 10.78; p <0.005), but no significant effect of dose (F[2,31] = 0.59; ns), and no significant dose x microinjection interaction (F [2,31] = 2.97; ns). However, a trend is observed for this interaction (p = 0.06). Note that for the dose factor, the shape of the DAergic response curves, opposite for the PBS and TTX groups, might explain the lack of statistical significance.

Regarding the time course of DAergic responses, the general ANOVA performed on the 37 animals during the first 60 min post-injection shows a highly significant effect of time (F[29,899] = 8.98; p <0.00001), a highly significant time x dose interaction (F[58,899] = 2.71; p <0.00001), a significant time x microinjection interaction (F[29,899] = 2.53; p <0.0001), as well as a significant time x dose x microinjection interaction (F[58,899] = 1.56; p <0.01). Similarly, for the 120 min post-injection, the ANOVA performed on the 37 animals reveals a highly significant effect of time (F [59.1829] = 7.29; p <0.00001), a highly significant time x dose interaction (F [118,1829] = 2.14; p <0.00001), a highly significant time x microinjection interaction (F [59,1829] = 2.65; p <0.00001), and a significant time x dose x microinjection interaction (F[118,1829] = 1.66; p <0.0001).

To summarize, the DAergic variations in the shell part of the left Nacc following the administration of MK-801 are significantly dependent on the neonatal microinjection in the left VSub(PBS or TTX). Furthermore, the time-course of DAergic responses appears to depend on the dose administered (NaCl 0.9%; MK-801 0.1 mg/kg; MK-801 0.2 mg/kg), the neonatal microinjection, and the interaction between these 2 factors. The interactions are significant during both the first 60 min post-injection period and the 120 min post-injection period.

## 4. DISCUSSION

The present study falls within the framework of animal modeling of certain aspects of the pathophysiology of schizophrenia. Its aim was to characterize and clarify the consequences of the neonatal functional inactivation of left VSub at PND8, in adult rats, after sc administration of a non-competitive NMDA receptor antagonist, exhibiting high selectivity and affinity for these receptors, MK-801 (dizocilpine) (Reynolds et al., 1987; Ogden and Traynelis, 2011). Two indices were used: locomotor activity and DAergic responses in the dorsomedial *shell* part of the Nacc. It is important to remember that locomotor activity is of particular interest in the context of animal modeling for schizophrenia because of the parallels that can be drawn between the emergence of hyperlocomotion in animals and psychotic symptoms in humans (Van den Busse, 2010). Also, the dorsomedial *shell* part of the Nacc appears to be the common target structure for the different types of anti-psychotics, all of which have the property of blocking D2-type DAergic receptors (Deutch et al., 1992, Merchant and Dorsa, 1993; Mo et al., 2005; Jennings et al., 2006; Natesan et al., 2006; Collins et al., 2014). The dorsomedial *shell* part of the Nacc therefore appears to be key for understanding the pathophysiology of schizophrenia, adding to the interest of this study. The results obtained show, first, that after administration of MK-801, locomotor hyperactivity is observed in both groups, the control (PBS) group and the experimental (TTX) group, and that such hyperactivity is significantly higher for the weakest dose of MK-801 (0.1 mg/kg sc) in TTX animals. Second, as regards DAergic responses, animals microinjected with PBS in the VSub display a rapid decrease in the DAergic response in the dorsomedial *shell* of the Nacc after sc injection of MK-801, whereas animals microinjected with TTX show a marked increase in the DAergic response in this striatal subregion.

### 1. Locomotor activity

The results obtained show first there is no difference between the PBS and TTX groups as regards spontaneous locomotor activity preceding the injection of 0.9% NaCl or MK-801. This suggests there are no basal changes in locomotor activity in TTX animals. The sc injection of 0.9% NaCl produces a slight transient increase in locomotor activity during the 10 min post-injection for the two PBS or TTX groups, but no notable variation or difference between the two groups is observed for the 2 hours post-injection. By contrast, injection of MK-801 gives rise to dose-dependent locomotor hyperactivity for both PBS and TTX groups. For the PBS group, the results are comparable to those reported by other authors (Pouvreau et al., 2016; Tagliabue et al., 2017; Wegener et al., 2011) showing, for identical doses of MK-801, an increase in locomotor activity with a distinctly greater effect for the highest dose of MK-801 (0.2 mg/kg sc) from the 20 min post-injection, an effect which begins to decline after 70 min. In the TTX group a very marked increase, peaking after 30 min, was observed following injection of the lowest dose of MK-801 (0.1 mg / kg sc). For this dose, locomotor hyperactivity not only peaks earlier than for the PBS group but is also significantly different during the 60 min post-injection.

The originality of the present study is underscored by the fact that, to the best of our knowledge, the effects of MK-801 after neonatal VSub blockade by TTX have never been reported. However, it is interesting to compare the results with those described after neonatal functional inactivation, by TTX at PND7, of the ventral hippocampus, a major afferent in the VSub (Lipska et al., 2002). In the latter study, exacerbated hyperlocomotion is obtained in TTX animals compared to PBS animals after administration of MK-801, but only for the highest dose of 0.2 mg/kg sc, in contrast to the results of the present study. The explication for the diverging results between these two studies might be the age difference of the animals tested, i.e., PND56 in the Lipska’s study (2002) as opposed to PND77 in the present study, and/or the difference in the TTX microinjection site, which was either the ventral hippocampus or the VSub. Whatever the case may be, the greater locomotor hyperactivity of the rats having undergone neonatal functional inactivation of the VSub with the MK-801 dose of 0.1 mg/kg sc is compatible with an increase in the sensitivity of the NMDA receptors and/or the number of such receptors in TTX microinjected animals. It is noteworthy in this respect that after neonatal TTX inactivation of the prefrontal cortex (PFC), which receives afferents from the VSub (Thierry et al., 2000), a significant difference in hyperlocomotion was observed only with the higher dose of 0.2mg/kg sc (Pouvreau et al., 2016; Tagliabue et al., 2017), the explanation perhaps being that inactivation of the VSub has a greater impact than that of the PFC.

### 2. DAergic variations in the *shell* part of the Nacc

The injection of 0.9% NaCl did not induce any different or marked changes in DA levels for the animals in either the PBS or TTX groups. In contrast, the administration of MK-801 resulted in divergent variations in both PBS and TTX animals. Injection of the NMDA antagonist is followed, in PBS animals, first by a dose-dependent drop in the DAergic signal and then by a rebound upto and above basal values, whereas in TTX animals a rapid and sustained increase in DAergic signal is observed. As far as we know, the effects of MK-801 on extracellular DA levels in the dorsomedial *shell* part of the Nacc after neonatal inactivation of the VSub by TTX have never been described in the literature, making it difficult to draw comparisons with other studies. It is to be pointed out, however, that the divergent profile of DAergic TTX/PBS responses is similar, albeit not identical, to that obtained after TTX functional blockade of the PFC at PND8 (Pouvreau et al., 2016).

#### 2.1. Animals microinjected with PBS in the VSub at PND8

It might be possible to explain the data obtained in the present study based on the scheme already proposed and discussed by Pouvreau et al. (2016) for interpreting the DAergic variations observed in the dorsomedial *shell* of the Nacc after administering MK-801 in adults animals neonatally microinjected with PBS in the PFC. Basically, according to this interpretation scheme it might be the case in the present study concerning the VSub that opposite influences come into competition with each other after administration of MK-801: first, direct excitatory pathways involving NMDA receptors situated presynaptically on DAergic terminals in the *shell* part of the Nacc, whose blockade by MK-801 results in a reduction in extracellular DA release, and, second, indirect inhibitory pathways involving NMDA receptors located on GABAergic interneurons, whose inhibition by MK-801 action would cause extracellular DA release to rise (see Pouvreau et al., 2016, for details). The data obtained in the present study suggest the initial involvement of the excitatory pathways.

#### 2.2 Animals microinjected with TTX in the VSub at PND8

For animals that underwent neonatal functional inactivation of the VSub with TTX at PND8, a dose-dependent increase in DAergic signal was observed following administration of MK-801, unlike the DAergic decreases noted in PBS animals. Taking account of the aforementioned interpretation scheme devised by Pouvreau et al. (2016) to explain the results obtained in animals microinjected with PBS in the VSub, it seems possible to suggest that the specific response obtained in animals in the TTX group after administration of MK-801 might be due to preponderant involvement of indirect inhibitory pathways. In these animals the involvement of the excitatory pathways would be greatly diminished without being completely abolished as suggested by the absence of any increase in the DAergic signal during the 10–15 min post-injection. Regarding more specifically the involvement of indirect inhibitory pathways, the anatomical data show that the VSub sends direct glutamatergic projections to the dorsomedial *shell* part of the Nacc which at this level seems to innervate mainly neurons forming part of local circuits, possibly GABAergic (Meredith and Totterdel; 1999; French and Totterdell, 2004). Since, to the best of our knowledge, the VSub does not directly innervate the ventral tegmental area (VTA), the region of origin of DAergic neurons (Phillipson, 1979; Geisler et al., 2007), the simplest hypothesis would be that neonatal inactivation of the VSub by TTX results in an alteration of connectivity between the VSub and the dorsomedial Nacc *shell*, involving the NMDA receptors located in a postsynaptic position on the *shell* GABAergic interneurons (see Pouvreau et al., 2016).

However, it is worth mentioning that it has been proposed that the VSub influences DA levels in the Nacc via a polysynaptic indirect pathway involving projections from the VSub to the Nacc, and then projections from the Nacc to the VTA via the ventral pallidum (Lisman and Grace, 2005). First of all, a partial reduction in the number of spontaneously active VTA DAergic neurons was observed after TTX microinjected in the VSub, which is consistent with previous report in *adult* animals in *acute* preparation (Louilot and Le Moal, 1994). Conversely, NMDA stimulation of the VSub was followed by a dose-dependent increase in active VTA DAergic neurons. It appears unlikely, however, that the aforementioned functional polysynaptic pathway was involved in the DA variations we observed, insofar as the NMDA antagonist kynurenic acid microinjected locally in the *shell* part of the Nacc has no effect or inhibits the enhancement of VTA DAergic activity induced by VSub stimulation (Lisman and Grace, 2005). If the polysynaptic Nacc-VTA pathway was involved in the present study we would expect no change or a reduction in extracellular DA levels after MK-801 in TTX animals microinjected at PND8, whereas marked increases were observed (present study).

A second, parallel VSub-VTA indirect pathway involving glutamatergic projections from the VSub to the Bed nucleus of the stria terminalis (BNST) and from the BNST to the VTA has been reported by Glangetas et al. (2015). Microinjection of glutamatergic antagonists in the BNST (including the NMDA antagonist AP5) inhibit the excitatory response of VTA DAergic neurons induced by VSub stimulation. As discussed above for the Nacc-VTA pathway, if the BNST-VTA pathway was involved in the variations observed in the present study after MK-801 administration, no change or a decrease in DA levels would be expected in the *shell* part of the Nacc in animals neonatally microinjected with TTX, whereas on the contrary significant increases were observed.

Another, non-exhaustive possibility would be the contribution of an indirect VSub-Nacc pathway through the PFC. Indeed, projections from the VSub to the PFC (prelimbic/infralimbic regions) (Thierry et al., 2000) and efferents from these PFC regions to the *shell* part of the Nacc have been well described, with a convergence of PFC and VSub projections on the same neurons in the *shell* part of the Nacc (Sesack and Grace, 2009). However, inasmuch as the PFC has been found to exert a complex permissive role on direct VSub-Nacc afferents, the putative contribution of a multisynaptic VSub-PFC-Nacc pathway would not be independent, but rather cooperative with that of the direct VSub-Nacc pathway. Interestingly, DAergic variations induced by MK-801 administration as observed in the present study seem faster and more marked than those previously observed after neonatal TTX blockade of the PFC (Pouvreau et al., 2016).

To summarize, the most parsimonious hypothesis explaining the increased DAergic responses consecutive to MK-801 administration in microinjected TTX animals at PND8 (i.e. a critical point in the neurodevelopmental period) is the involvement of abnormal functioning of NMDA receptors present on GABAergic interneurons situated in the dorsomedial *shell* part of the Nacc. In this context, it is important to remember that NMDA receptors are multimeric assemblies of NR1, NR2 and NR3 subunits, with most central NMDA receptors being NR1/NR2 assemblies (Lau and Zukin, 2007), and that NMDA receptors including a NR3 subunit (NR3A or NR3B) are insensitive to open-channel NMDA receptor blocking agents such as MK-801 (Perez-Otano et al., 2016). Typically, NMDA receptors comprise two NR1-type subunits and two NR2-type subunits. The NR1 subunit is generally considered indispensable for forming a functional NMDA receptor, whereas the identity of the NR2 subunit (NR2A, NR2B, NR2C and NR2D) determines many of the properties of NMDA receptors, particularly pharmacological properties. During postnatal development the expression and number of NR2A subunits/NR2B subunits go up, resulting in changes in the composition of NMDA receptors. Furthermore, interruption of the electrical neuronal activity by TTX induces an increase in the expression of the two types of NR2 subunits and in the synaptic number of NMDA receptors (Lau and Zukin, 2007). It is also noteworthy that synaptic NMDA receptors are highly concentrated in the post-synaptic densities (PSD) of excitatory synapses, where they are associated with several scaffolding and signaling proteins (Sheng and Kim, 2011, Kaizuka and Takumi, 2018), and that, furthermore, TTX blockade triggers a change in PSD conformation (Glebov et al., 2016). So it is tempting to suggest that a conformational change in the PSD at the level of the dorsomedial *shell* is also involved in the present results. It is of interest here to note that recent post-mortem studies described a reduction in the size of the PSD at the level of asymmetric synapses in the *shell* subregion of the Nacc in patients with schizophrenia (McCollum et al., 2015).

## 5. CONCLUSION

To conclude, the present results support the hypothesis that neonatal VSub functional inactivation produces an increase in both locomotor and DAergic responses in the dorsomedial *shell* part of the Nacc following administration of MK-801 in adult rats. They further suggest that an alteration in glutamatergic transmission involving NMDA receptors could be responsible for dysregulation in these responses, particularly the increase in the *shell* DAergic release, although the specific mechanisms behind changes in these responses after neonatal inactivation merit clarification.

Finally, to further validate our approach dealing with the VSub in the context of animal modeling of schizophrenia (Meyer et al., 2009, Meyer and Louilot, 2011, 2014; Saoud et al., 2020) it would be interesting, after neonatal functional blockade of the VSub, to investigate possible changes in PSD-associated proteins in the *shell* part of the Nacc. Indeed, it has recently been suggested that dysregulation of PSD-95 protein expression is involved in the pathophysiology of schizophrenia (Funk et al., 2017).

## Acknowledgements

The authors are grateful to Solène Begaudeau, Mi-Hae Hanen, Caroline Lahogue and Laurine Stroh for their helpful assistance.

## References

Briend, F., Nelson, E.A., Maximo, O., Armstrong, W.P., Kraguljac, N.V., Lahti, A.C., 2020. Hippocampal glutamate and hippocampus subfield volumes in antipsychotic-naive first episode psychosis subjects and relationships to duration of untreated psychosis. Transl. Psychiatry. 10, 137.

Collins, C.M., Wood, M.D., and Elliott, J.M., 2014. Chronic administration of haloperidol and clozapine induces differential effects on the expression of Arc and c-Fos in rat brain. Journal of Psychopharmacology. 28, 947–954.

Coyle, J.T., 2012. NMDA Receptor and Schizophrenia: A Brief History. Schizophr. Bull. 38, 920–926.

Deutch, A.Y., Lee, M.C., Iadarola, M.J., 1992. Regionally specific effects of atypical antipsychotic drugs on striatal Fos expression: The nucleus accumbens shell as a locus of antipsychotic action. Molecular and Cellular Neuroscience. 3, 332–341.

Field, K.J., White, W.J., Lang, C.M., 1993. Anaesthetic effects of chloral hydrate, pentobarbitone and urethane in adult male rats. Lab. Anim. 27, 258–269.

Fornito, A., Zalesky, A., Pantelis, C., Bullmore, E.T., 2012. Schizophrenia, neuroimaging and connectomics. NeuroImage. 62, 2296–2314.

French, S.J., Totterdell, S., 2004. Quantification of morphological differences in boutons from different afferent populations to the nucleus accumbens, Brain Res. 1007, 167–177.

Friston, K.J., 1996. Theoretical Neurobiology and Schizophrenia. Br. Med. Bull. 52, 644–655.

Funk, A.J., Mielnik, C.A., Koene, R., Newburn, E., Ramsey, A.J., Lipska, B.K., McCullumsmith, R.E., 2017. Postsynaptic Density-95 isoform abnormalities in schizophrenia. Schizophr. Bull. 43, 891–899.

Geisler, S., Derst, C., Veh R.W., Zahm D.S., 2007. Glutamatergic afferents of the ventral tegmental area in the rat. J. Neurosci. 27, 5730–5743.

Giezendanner, S., Walther, S., Razavi, N., Swam1, C.V., Fisler, M.S., Soravia, L.M., Andreotti, J., Schwab, S., Jann, K., Wiest, R., Horn, H., Müller, T.J., Dierks, T., Federspiel, A., 2013. Alterations of White Matter Integrity Related to the Season of Birth in Schizophrenia: A DTI Study. PLos One, 8, e75508.

Glangetas, C., Fois, G.R., Jalabert, M., Lecca, S., Valentinova, K., Meye, F.J., Diana, M., Faure, P., Mameli, M.,, Caille, S., Georges, F., 2015. Ventral Subiculum Stimulation Promotes Persistent Hyperactivity of Dopamine Neurons and Facilitates Behavioral Effects of Cocaine. Cell Rep. 13, 2287–2296.

Glebov, O.O., Cox, S., Humphreys, L., Burrone, J. 2016. Neuronal activity controls transsynaptic geometry. Sci Rep. 6, 22703.

González-Mora, J.L., Pedro Salazar, P., Martín, M., Mas, M., 2017. Monitoring Extracellular Molecules in Neuroscience by In Vivo Electrochemistry: Methodological Considerations and Biological Applications. In: Athineos Philippu (ed.), In Vivo Neuropharmacology and Neurophysiology, Neuromethods, Springer Science+Business Media, New York, vol. 121, 181–206.

Itil, T., Keskiner, A., Kiremitci, N., Holden, J.M., 1967. Effect of phencyclidine in chronic schizophrenics. Can. Psychiatr. Assoc. J. 12, 209–212.

Jennings, C.A., Cluderay, J.E., Gartlon, J., Cilia, J., Lloyd, A., Jones, D.N., Southam, E., 2006. The effects of ziprasidone on regional c-Fos expression in the rat forebrain. Psychopharmacology (Berl) 184, 13–20.

Kaar, S.J., Natesan, S., McCutcheon, R., Howes, O.D., 2020. Antipsychotics: Mechanisms underlying clinical response and side-effects and novel treatment approaches based on pathophysiology. Neuropharmacology. 172, 107704.

Kaizuka, T., Takumi, T., 2018. Postsynaptic density proteins and their involvement in neurodevelopmental disorders. J. Biochem. 163, 447–455.

Karlsgodt, K.H., Sun, D., Jimenez, A.M., Lutkenhoff, E.S., Willhite, R., van Erp, T.G., Cannon, T.D., 2008. Developmental disruptions in neural connectivity in the pathophysiology of schizophrenia. Dev. Psychopathol. 20, 1297–1327.

Lau, C.G., Zukin, R.S., 2007. NMDA receptor trafficking in synaptic plasticity and neuropsychiatric disorders. Nat. Rev. Neurosci. 8, 413–426.

Leucht, S., Leucht, C., Huhn, M., Chaimani, A., Mavridis, D., Helfer, B., Samara, M., Rabaioli, M., Bächer, S., Cipriani, A., Geddes, J.R., Salanti, G., Davis, J.M., 2017. Trials in Acute Schizophrenia: Systematic Review, Bayesian Meta-Analysis, and Meta-Regression of Efficacy Predictors. Am. J. Psychiatry. 174, 927–942.

Lipska, B.K., Halim, N.D., Segal, P.N., Weinberger, D.R., 2002. Effects of reversible inactivation of the neonatal ventral hippocampus on behavior in the adult rat. J. Neurosci. 22, 2835–2842.

Lisman, J.E., Grace A.A., 2005. The hippocampal-VTA loop: controlling the entry of information into long-term memory. Neuron. 46, 703–713.

Louilot, A., Le Moal, M., 1994. Lateralized interdependence between limbicotemporal and ventrostriatal dopaminergic transmission Neuroscience. 59, 495–500.

Louilot, A., Serrano, A., D’Angio, M., 1987. A novel carbon fiber implantation assembly for cerebral voltammetric measurements in freely moving rats. Physiol. Behav. 41, 227–231.

McCollum, L.A., Walker, C.K., Roche, J.K., Roberts, R.C., 2015. Elevated Excitatory Input to the Nucleus Accumbens in Schizophrenia: A Postmortem Ultrastructural Study. Schizophr. Bull. 41, 1123–1132.

McCutcheon, R., Beck, K., Jauhar, S., Howes, O.D., 2018. Defining the locus of dopaminergic dysfunction in schizophrenia: A meta-analysis and test of the mesolimbic hypothesis. Schizophr. Bull. 44, 1301–1311.

McGlashan, T.H., Hoffman, R.E., 2000. Schizophrenia as a disorder of developmentally reduced synaptic connectivity. Arch. Gen. Psychiatry. 57, 637–648.

Merchant, K.M., Dorsa, D.M., 1993. Differential induction of neurotensin and c-fos gene expression by typical versus atypical antipsychotics. Proc. Natl. Acad. Sci. USA 90, 3447–3451.

Meredith, G.E., Totterdell,S., 1999. Microcircuits in nucleus accumbens’ shell and core involved in cognition and reward. Psychobiology. 27, 165–186.

Meyer, F., Louilot, A., 2011. Latent inhibition-related dopaminergic responses in the nucleus accumbens are disrupted following neonatal transient inactivation of the ventral subiculum. Neuropsychopharmacology. 36, 1421–1432.

Meyer, F., Louilot, A., 2012. Early prefrontal functional blockade in rats results in schizophrenia-related anomalies in behavior and dopamine. Neuropsychopharmacology. 37, 2233–2243.

Meyer, F., Louilot, A., 2014. Consequences at adulthood of transient inactivation of the parahippocampal and prefrontal regions during early development: new insights from a disconnection animal model for schizophrenia. Front. Behav. Neurosci. 8, 118.

Meyer, F., Peterschmitt, Y., Louilot, A., 2009. Postnatal functional inactivation of the entorhinal cortex or ventral subiculum has different consequences for latent inhibition-related striatal dopaminergic responses in adult rats. Eur. J. Neurosci. 29, 2035–2048.

Mo, Y.Q., Jin X.L., Chen, Y.T., Jin, G.Z., Shi, W.X., 2005, Effects of l-Stepholidine on Forebrain Fos Expression: Comparison with Clozapine and Haloperidol. Neuropsychopharmacology. 30, 261–267.

Moyer, C.E., Shelton, M.A., Sweet, R.A., 2015. Dendritic spine alterations in schizophrenia. Neurosci Lett. 601, 46–53.

Natesan, S., Reckless, G.E., Nobrega, J.N. Fletcher, P.J., Kapur, S., 2006. Dissociation between In Vivo Occupancy and Functional Antagonism of Dopamine D2 Receptors: Comparing Aripiprazole to Other Antipsychotics in Animal Models. Neuropsychopharmacology. 31, 1854–1863.

Nikolaus, S., Mamlins, E., Hautzel, H., Müller H.-W., 2019. Acute anxiety disorder, major depressive disorder, bipolar disorder and schizophrenia are related to different patterns of nigrostriatal and mesolimbic dopamine dysfunction. Rev. Neurosci. 30, 381–426.

Ogden, K.K., Traynelis, S.E., 2011. New advances in NMDA receptor pharmacology Trends Pharmacol. Sci. 32, 726–733.

Paxinos, G., Watson, C., 2014. The Rat Brain in Stereotaxic Coordinates, 7th edn., Academic Press, San Diego.

Pérez-Otaño, I., Larsen, R.S., Wesseling, J.F., 2016. Emerging roles of GluN3-containing NMDA receptors in the CNS. Nat Rev. Neurosci.17, 623–635.

Phillipson, O.T., 1979. Afferent projections to the ventral tegmental area of Tsai and interfascicular nucleus: a horseradish peroxidase study in the rat. J. Comp. Neurol. 187, 117–143.

Pouvreau, T., Tagliabue, E., Usun, Y., Eybrard, S., Meyer, F., Louilot, A., 2016. Neonatal Prefrontal Inactivation Results in Reversed Dopaminergic Responses in the Shell Subregion of the Nucleus Accumbens to NMDA Antagonists. ACS. Chem. Neurosci. 7, 964–971.

Reynolds, I.J., Murphy, S.N., Miller, R.J., 1987. 3H-labeled MK-801 binding to the excitatory amino acid receptor complex from rat brain is enhanced by glycine. Proc. Natl. Acad. Sci. USA. 84, 7744–7748.

Rioux, L., Nissanov, J., Lauber, K., Bilker, W.B., Arnold, S.E., 2003. Distribution of microtubule-associated protein MAP2-immunoreactive interstitial neurons in the parahippocampal white matter in subjects with schizophrenia. Am. J. Psychiatry. 160, 149–155.

Rosenthal, R., Rosnow, R.L., Rubin, D.B., 2000. Contrasts and effect sizes in behavioral research: a correlational approach. Cambridge University Press, New York.

Saoud, H., De Beus, D., Eybrard, S., Louilot, A., 2020. Postnatal functional inactivation of the ventral subiculum enhances dopaminergic responses in the core part of the nucleus accumbens following ketamine injection in adult rats. Neurochem Int. 137, 104736.

Seeman, P., 1987. Dopamine receptors and the dopamine hypothesis of schizophrenia. Synapse. 1, 133–152.

Sesack, S.R., Grace, A.A., 2009. Cortico-Basal Ganglia Reward Network: Microcircuitry. Neuropsychopharmacology, 35, 27–47.

Sheng, M., Kim, E., 2011. The postsynaptic organization of synapses. Cold Spring Harb Perspect. Biol. 3, a005678.

Tagliabue, E., Pouvreau, T., Eybrard, S., Meyer, F., Louilot, A., 2017. Dopaminergic responses in the core part of the nucleus accumbens to subcutaneous MK801 administration are increased following postnatal transient blockade of the prefrontal cortex. Behav. Brain. Res. 335, 191–198.

Talamini, L.M., Meeter, M., Elvevåg, B., Murre, J.M.J., Terry E Goldberg, T.E., 2005. Reduced Parahippocampal Connectivity Produces Schizophrenia-like Memory Deficits in Simulated Neural Circuits With Reduced Parahippocampal Connectivity. Arch. Gen. Psychiatry. 62, 485–493.

Tandon, R., Gaebel, W., Barch, D.M., Bustillo, J., Gur, R.E., Heckers, S., Malaspina, D., Owen, M.J., Schultz, S., Tsuang, M., Van Os, J., Carpenter, W., 2013. Definition and description of schizophrenia in the DSM-5. Schizophr. Res. 150, 3–10.

Thierry, A.-M., Gioanni, Y., Dégénétais, E., Glowinski, J., 2000. Hippocampo-Prefrontal Cortex Pathway: Anatomical and Electrophysiological Characteristics. Hippocampus. 10, 411–419.

Usun, Y., Eybrard, S., Meyer, F., Louilot, A., 2013. Ketamine increases striatal dopamine release and hyperlocomotion in adult rats after postnatal functional blockade of the prefrontal cortex. Behav. Brain Res. 256, 229–237.

Van den Buuse, M., 2010. Modeling the Positive Symptoms of Schizophrenia in Genetically Modified Mice: Pharmacology and Methodology Aspects. Schizophr. Bull. 36, 246–270.

Van Rossum, J.M., 1966. The significance of dopamine-receptor blockade for the mechanism of action of neuroleptic drugs. Arch. Int. Pharmacodyn. Ther., 160, 492–494.

Weinberger, D.R., 2017. Future of Days Past: Neurodevelopment and Schizophrenia. Schizophr. Bull. 43, 1164–1168.

Wegener, N., Nagel, J., Gross, R., Chambon, C., Greco, S., Pietraszek, M., Gravius, A., and Danysz, W., 2011. Evaluation of brain pharmacokinetics of (+)MK-801 in relation to behaviour. Neurosci. Lett. 503, 68–72.

